# *Xanthomonas indica* sp. nov., a non-pathogenic bacterium isolated from healthy rice *(Oryza sativa)* seeds from India

**DOI:** 10.1101/2022.03.16.484583

**Authors:** Rekha Rana, Vishnu Narayanan Madhavan, Tanu Saroha, Kanika Bansal, Amandeep Kaur, Ramesh V. Sonti, Hitendra K. Patel, Prabhu B. Patil

## Abstract

Two yellow pigmented bacterial strains were isolated from healthy rice seeds. The strains designated as PPL560^T^ and PPL568 were identified as members of genus *Xanthomonas* based on analysis of biochemical and 16S rRNA gene sequence retrieved from whole genome sequence. Isolates formed a distinct monophyletic lineage with *X. sontii* and *X. sacchari* as the closest relatives in the phylogenetic tree based on core gene content shared by reported species in genus *Xanthomonas*. Pairwise ortho Average Nucleotide Identity and digital DNA-DNA hybridisation values calculated against other species of *Xanthomonas* were below their respective cut-offs. *In planta* studies revealed that PPL560^T^ and PPL568 are non-pathogenic to rice plants upon leaf clip inoculation. Absence of type III secretion system related genes and effectors further supported their non-pathogenic status. Herein, we propose *Xanthomonas indica* sp. nov. as novel species of genus *Xanthomonas* with PPL560^T^=MTCC13185 as its type strain and PPL568 as another constituent member.

## Introduction

*Xanthomonas* (Dowson, 1939) is a group of large number of plant pathogenic bacteria. The member species and pathovars infect a wide range of host plants in host and tissue specific manner (Young, Dye, Bradbury, Panagopoulos, & Robbs, 1978). In the beginning, species of this complex genus were categorized using the ‘new host-new species’ concept based on the assumption that pathogens infecting different hosts must be different in their biochemical properties (Luc Vauterin, Rademaker, & Swings, 2000). Vauterin et al. (1995), grouped xanthomonads in 20 different species based on DNA-DNA hybridization analysis (L Vauterin, Hoste, Kersters, & Swings, 1995), which was gold standard for species characterization of genus *Xanthomonas* in consequent years (J. B. Jones, Lacy, Bouzar, Stall, & Schaad, 2004; Schaad et al., 2005; Trébaol et al., 2000). There are 31 validly published species in the genus that infect around 400 plants including both monocots and dicots (Parte, Carbasse, Meier-Kolthoff, Reimer, & Göker, 2020; Ryan et al., 2011). With the technological breakthroughs and reduced costs of whole genome sequencing data, species delineation has become an easier and reproducible task. Average nucleotide identity (ANI) and digital DNA-DNA hybridisation (dDDH) calculation between two genome sequences and construction of core gene phylogenies have emerged as useful tools for novel species characterization (Bansal et al., 2021; López et al., 2018; Martins et al., 2020). A value of about 95%-96% ANI is considered equivalent to 70% dDDH/DDH value, which are accepted cut-offs for species delineation (Goris et al., 2007; Richter & Rosselló-Móra, 2009; Wayne et al., 1987). When combined with the biochemical analysis, genome based taxonomics is rapidly emerging as a powerful approach for the reclassification of existing strains as well as for the description of novel species in the genus *Xanthomonas* (da Gama et al., 2018; Martins et al., 2020).

Genus *Xanthomonas* largely comprises of pathogens, however increasingly *Xanthomonas* strains are being isolated from healthy plant parts (Di Ming, Schaad, & Roth, 1991; R. Jones et al., 1989; Luc Vauterin et al., 1996). These strains, when inoculated into their hosts fail to produce disease symptoms and most do not belong to any previously reported pathogenic species of genus *Xanthomonas* (Cottyn, Debode, Regalado, Mew, & Swings, 2009; Cottyn et al., 2001). These non-pathogenic strains are largely overlooked when compared to pathogenic ones for they lack economic importance. However, these strains need to be properly classified so their identification can be differentiated from that of pathogens for crop disease management, quarantine and regulation purposes. Moreover, horizontal gene transfer events among pathogens and non-pathogens from the same host plant might lead to the emergence of new much more deadly pathogens (Merda et al., 2016). These non-pathogenic *Xanthomonas* might as well be related to biocontrol activities against host plant diseases (Li et al., 2020). Keeping the above points in mind, it is imperative to understand their ecology and relationship with pathogenic strains.

Rice is a staple food crop and non-pathogenic *Xanthomonas* strains are repeatedly found associated with healthy rice plants (Cottyn et al., 2009; Cottyn et al., 2001; Di Ming et al., 1991). Recently, two new species, *X. maliensis* and *Xanthomonas sontii*, with non-pathogenic strains were reported to be associated with rice plants (Bansal et al., 2021; Triplett et al., 2015). In the present study, we report two strains of a potential novel non-pathogenic species isolated from healthy rice seeds. Our biochemical and genome comparison analyses revealed them to be distinct from previously reported *Xanthomonas* species from rice and other plants. Pathogenicity assay on rice plants indicated that these strains are non-pathogenic in nature. Here, we propose *Xanthomonas indica* sp. nov. with PPL560^T^ as type strain and PPL568 as another diverse member of this novel species.

## Bacterial isolation

Healthy rice seeds belonging to USHA-44 cultivar were obtained directly from farmers’ fields in Ropar, Punjab (India) in 2020 during the harvest season. For bacterial isolation we followed the protocol mentioned in the literature (Cottyn et al., 2001; Midha et al., 2016). Briefly, rice seeds were separated from husk using sterile forceps. 5g rice seeds were partially crushed using autoclaved pestle mortar and suspended in 50 ml of 0.85% NaCl solution. The solution was incubated at 28°C with constant shaking at 180 rpm for 2 hours. The solution was serially diluted up to 10^-4^ in 0.85% NaCl and plated in duplicates on 4 different media, Peptone Sucrose Agar (PSA), Nutrient Agar (NA), Glucose Yeast extract Calcium carbonate Agar (GYCA), and Tryptone Soya Agar (TSA), each containing 0.01% cycloheximide. These plates were incubated at 28°C for a time period of a week with continuous growth monitoring. Plates only with 0.85% saline and without saline were used as control to check contamination. All the mucoid yellow coloured single colonies were further streaked to obtain pure cultures on NA as well as PSA. The strains were stored in 15% glycerol at −80°C. The strains, PPL560^T^ and PPL568 are deposited to national as well as international microbial culture collections with accession numbers, PPL560^T^=MTCC13185 and PPL568=MTCC13186.

## Taxonogenomics of PPL560^T^ and PPL568

Genomic DNA was extracted from overnight cultures of PPL560^T^ and PPL568 grown in PS (1% peptone and 1% sucrose) media using Quick-DNA^TM^ Fungal/Bacterial Miniprep Kit (Zymo Research, USA). Quantitative and qualitative analysis of genomic DNA was carried out using Qubit 4 fluorometer (Invitrogen; Thermo Fisher Scientific) and Nanodrop 1000 (Thermo Fisher Scientific). Genomic DNA was sequenced using Illumina Novaseq platform facility (MedGenome, Hyderabad). Quality assessment of the raw reads was done using FastQC version 0.11.9 (https://qubeshub.org/resources/fastqc). Sequenced paired end reads were assembled using SPAdes version 3.13.0 (Prjibelski, Antipov, Meleshko, Lapidus, & Korobeynikov, 2020). The genome quality was assessed using Quast version 5.0.2 (Gurevich, Saveliev, Vyahhi, & Tesler, 2013). The size and GC content of the genomes are approximately 4.7 Mb and 69.4%. The whole genomes of PPL560^T^ and PPL568 consists of 29 and 18 contigs with N50 values being in the range of 260kb to 440kb. Their average genome coverage was 735x and 1,317x for PPL560^T^ and PPL568, respectively. Further, genomes were checked for completeness and contamination using CheckM version 1.1.3 (Parks, Imelfort, Skennerton, Hugenholtz, & Tyson, 2015). Detailed genome assembly and annotation statistics are given in **Table 1**. Complete 16S rRNA gene sequence retrieved from the whole genome showed the closest identity to *X. sontii* PPL1^T^ in EZBioCloud database (Yoon et al., 2017). Whole genome sequences of PPL560^T^ and PPL568 were submitted to NCBI with accession numbers JAKJPQ000000000 and JAKLXA000000000, and annotated using the NCBI’s (https://www.ncbi.nlm.nih.gov/genome/annotation_prok/) Prokaryotic Genome Annotation Pipeline (PGAP).

**Table 1:** Genome assembly and annotation statistics for PPL560^T^ and PPL568.

| Statistics | PPL560<sup>T</sup> | PPL568 |
| --- | --- | --- |
| Genome Size (Mb) | 4.79 | 4.74 |
| G+C content (%) | 69.47 | 69.40 |
| Genome Coverage (x) | 735 | 1317 |
| Contigs | 29 | 18 |
| N50 (bp) | 262,450 | 439,146 |
| CDS | 4,030 | 4,044 |
| Completeness/Contamination | 100/0.00 | 100/0.00 |
| RNAs | 65 | 68 |
| Pseudogenes | 31 | 26 |
| Accession number | JAKJPQ000000000 | JAKLXA000000000 |

For PPL560^T^ and PPL568, pairwise ortho Average Nucleotide Identity (orthoANI) and digital DNA-DNA hybridisation (dDDH) values were calculated against type strains of all the species of genus *Xanthomonas* (Parte et al., 2020; www.bacterio.net). The orthoANI value was calculated using oANI (Lee, Kim, Park, & Chun, 2016) implemented with USEARCH version 11.0.667 and dDDH values were calculated using formula 2 of Genome to Genome Distance Calculator version 3.0 (Meier-Kolthoff, Carbasse, Peinado-Olarte, & Göker, 2021). orthoANI and dDDH values of PPL560^T^ and PPL568 were ≤93.5% and <60% with other species of genus *Xanthomonas*, which is below species delineation cut-offs **(Table 2)**. The strains had *X. sacchari* and *X. sontii* as closest relatives with their orthoANI values being in the range of 93.2% to 93.5% while for the other *Xanthomonas* spp. the values were below 90%. PPL560^T^ and PPL568 had dDDH values above 50% only in the case of *X. sacchari* and *X. sontii*, all other species had values much below the cut off.

**Table 2:** Percent digital DNA-DNA Hybridization (dDDH), and Ortho Average Nucleotide identity (orthoANI) values between PPL560^T^ and PPL568 strains and type strains of *Xanthomonas* spp.

| S. No. | Strains | dDDH |  | ANI |  |
| --- | --- | --- | --- | --- | --- |
|  |  | PPL560 | PPL568 | PPL560 | PPL568 |
| 1. | PPL560 | 100 | 68.0 | 100 | 96.3 |
| 2. | PPL568 | 68.0 | 100 | 96.3 | 100 |
| 3. | *X. sacchari* CFBP4641<sup>T</sup> | 51.9 | 51.2 | 93.5 | 93.5 |
| 4. | *X. sontii* PPL1<sup>T</sup> | 51.0 | 50.4 | 93.3 | 93.2 |
| 5. | *X. hyacinthi* CFBP1156<sup>T</sup> | 33.7 | 33.4 | 87.7 | 87.7 |
| 6. | *X. translucens* DSM18974<sup>T</sup> | 32.9 | 32.9 | 87.2 | 87.2 |
| 7. | *X. theicola* CFBP4691<sup>T</sup> | 32.5 | 32.5 | 87.2 | 87.0 |
| 8. | *X. albilineans* CFBP2523<sup>T</sup> | 28.5 | 28.4 | 84.3 | 84.5 |
| 9. | *X. euvesicatoria* LMG27970<sup>T</sup> | 23.6 | 23.4 | 79.3 | 79.4 |
| 10. | *X. arboricola* CFBP2528<sup>T</sup> | 23.4 | 23.3 | 79.9 | 79.7 |
| 11. | *X. codiae* CFBP4690<sup>T</sup> | 23.4 | 23.4 | 80.0 | 80.1 |
| 12. | *X. euroxanthea* CPBF424<sup>T</sup> | 23.4 | 23.5 | 80.2 | 80.0 |
| 13. | *X. alfalfae* LMG495<sup>T</sup> | 23.3 | 23.2 | 79.4 | 79.5 |
| 14. | *X. cassava* CFBP4642<sup>T</sup> | 23.3 | 23.3 | 79.5 | 79.7 |
| 15. | *X. melonis* CFBP4644<sup>T</sup> | 23.3 | 23.3 | 79.6 | 79.9 |
| 16. | *X. floridensis* WHRI8848<sup>T</sup> | 23.2 | 23.2 | 79.6 | 79.5 |
| 17. | *X. campestris* ATCC33913<sup>T</sup> | 23.1 | 22.9 | 79.5 | 79.3 |
| 18. | *X. axonopodis* DSM3585<sup>T</sup> | 23.0 | 22.9 | 79.0 | 79.0 |
| 19. | *X. cucurbitae* CFBP2542<sup>T</sup> | 23.0 | 23.0 | 79.7 | 79.5 |
| 20. | *X. hortorum* WHRI7744<sup>T</sup> | 23.0 | 22.9 | 79.3 | 79.1 |
| 21. | *X. nasturtii* WHRI8853<sup>T</sup> | 23.0 | 22.9 | 79.5 | 79.2 |
| 22. | *X. perforans* DSM18975<sup>T</sup> | 23.0 | 22.9 | 79.4 | 79.4 |
| 23. | *X. phaseoli* CFBP412 | 23.0 | 23.0 | 79.6 | 79.2 |
| 24. | *X. pisi* DSM18956<sup>T</sup> | 23.0 | 23.0 | 79.5 | 79.3 |
| 25. | *X. dyei* CFBP7245<sup>T</sup> | 22.9 | 22.8 | 79.1 | 79.1 |
| 26. | *X. maliensis* LMG27592<sup>T</sup> | 22.9 | 22.9 | 79.5 | 79.7 |
| 27. | *X. oryzae* ICMP3125<sup>T</sup> | 22.9 | 22.8 | 79.0 | 79.0 |
| 28. | *X. prunicola* CFBP8353<sup>T</sup> | 22.9 | 23.0 | 78.9 | 78.9 |
| 29. | *X. citri* LMG9322<sup>T</sup> | 22.9 | 23.0 | 79.2 | 79.4 |
| 30. | *X. bromi* CFBP1976<sup>T</sup> | 22.8 | 22.7 | 79.2 | 79.0 |
| 31. | *X. vesicatoria* LMG911<sup>T</sup> | 22.7 | 22.7 | 79.0 | 78.9 |
| 32. | *X. vasicola* NCPPB2417<sup>T</sup> | 22.6 | 22.8 | 78.6 | 78.7 |
| 33. | *X. populi* CFBP1817<sup>T</sup> | 22.5 | 22.6 | 78.8 | 78.9 |
| 34. | *X. fragariae* PD885<sup>T</sup> | 22.3 | 22.2 | 78.6 | 78.6 |

## Phylogenetic Analysis

The core gene phylogeny was constructed using type strains of *Xanthomonas* spp. along with novel strains PPL560^T^ and PPL568, taking *S. maltophilia* ATCC13637^T^ as an outgroup. The catenated alignment of core genes was obtained using Roary v3.13.0 (Page et al., 2015) generated taking Prokka v1.46 (Seemann, 2014) annotated files. Initial tree was generated using PhyML v3.3 (Guindon et al., 2010) with default parameters. Further, ClonalFramML v1.12 (Didelot & Wilson, 2015) was used to detect recombination regions which were then masked using maskrc-svg (https://github.com/kwongj/maskrc-svg). This masked alignment was used to build recombination free maximum likelihood phylogeny using PhyML v3.3 with General Time Reversible (GTR) nucleotide substitution model and 1000 bootstrap replicates. The phylogenetic tree was annotated using iTOL v6 (Letunic & Bork, 2021).

Core gene phylogeny of PPL560^T^ and PPL568 with strains of other *Xanthomonas* spp. demonstrated that they belong to clade I of *Xanthomonas* species consisting of *X. albilineans, X. hyacinthi, X. theicola, X. translucens*, *X. sontii* and *X. sacchari* (Koebnik et al., 2021; Mafakheri et al., 2022). The novel strains formed a monophyletic clade in close association with *X. sontii* and *X. sacchari*. The rest of the *Xanthomonas* spp. were forming another major group and *X. maliensis* was separated from these two major groups **(Figure 1)**.

**Figure 1.**
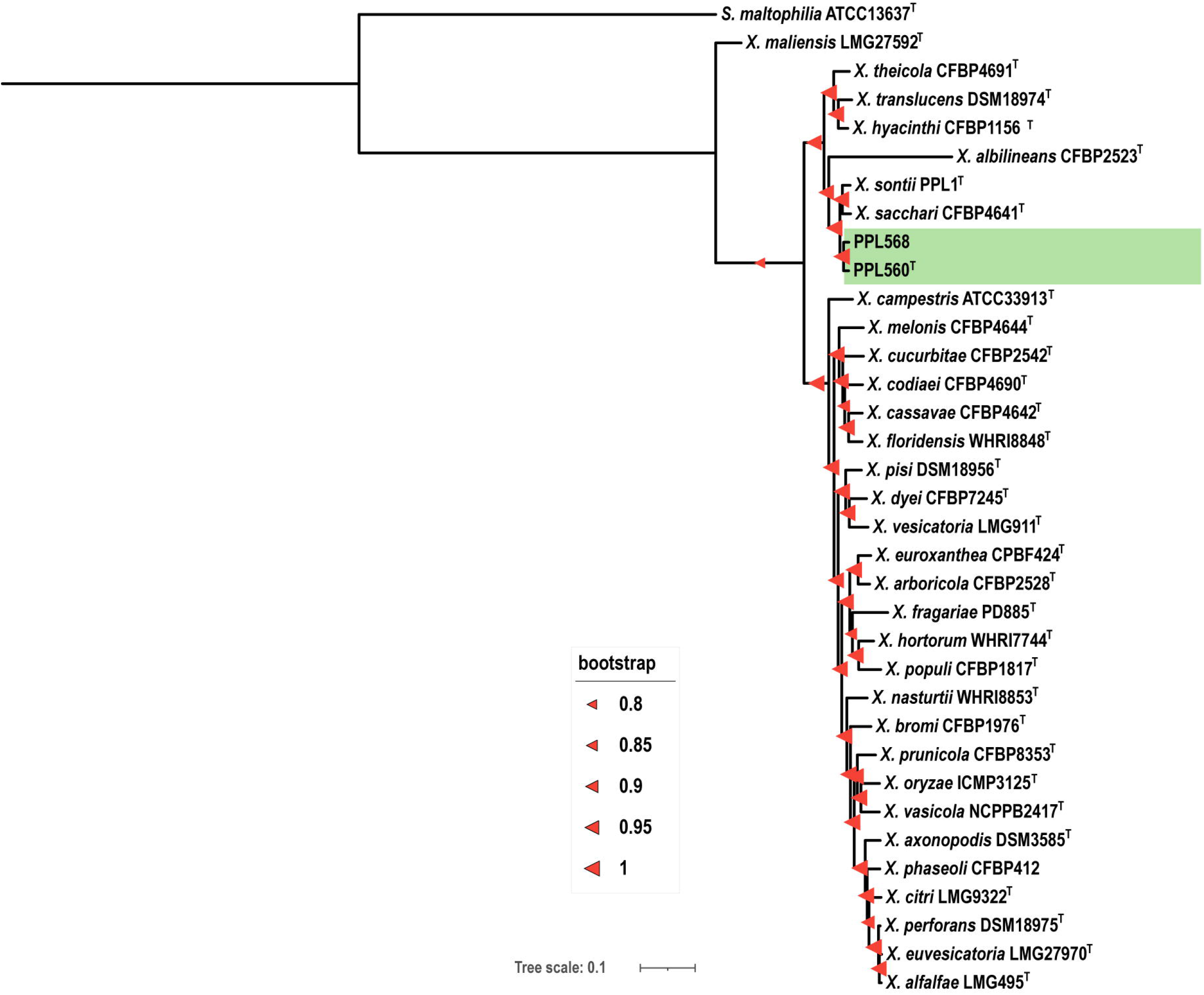
Recombination free phylogeny of novel strains and other *Xanthomonas* species. Maximum Likelihood phylogeny constructed using core gene content of PPL560^T^ and PPL568 with type strains of *Xanthomonas* species. The scale bar represents number of nucleotide substitutions per site. PPL560^T^ and PPL568 are highlighted in green colour. Red triangles at the nodes represent bootstrap values. Size of triangle is proportionate to the bootstrap value as depicted in the image.

## Phenotypic characterization

Biochemical characterization such as carbon utilization, acid production, and test for various enzymatic activities for novel strains, PPL560^T^ and PPL568 along with closely related *X. sontii* PPL1^T^ and *X. sacchari* NCPPB 4331^T^, was done with BIOLOG GN3 microplate system (Biolog Inc, USA) following protocol A as per manufacturer’s instructions. Briefly, bacterial strains were cultured on PSA at 28°C for 48h. Further, cultured colonies were suspended in IF-A fluid using sterile cotton swabs to obtain recommended turbidity range. The suspension was mixed properly using vortex and 100 μl was added to each well of BIOLOG GEN III microplate. The plates were incubated at 28°C and read using MicroStation 2 Reader at 24h and 48h, and results were interpreted using MicroLog 3/5.2.01 software. The tests were performed in triplicates and only +/− values are considered valid. Readings taken at 48h showed consistent results as compared to 24h, so taken up for further analysis. Biochemical profile was same for these four strains, complete biochemical profile of PPL560^T^ shown in **Supplementary Figure 1**. The strains were able to utilize Dextrin, D-Maltose, D-Trehalose, D-Cellobiose, Gentiobiose, Sucrose, D-Turanose, α-D-Lactose, D-Meliobiose, ß-Methyl-D-Glucoside, D-salicin, N-Acetyl-D-Glucosamine, α-D-Glucose, D-Mannose, D-Fructose, Gelatin, L-Alanine, L-Aspartic acid, L-Serine, L-Glutamic Acid, Pectin, Methyl pyruvate, L-Lactic acid, Citric acid, L-Malic acid, Bromo-succinic acid, Tween 40, Propionic acid, and Acetic acid. Strains were resistant to rifampicin SV, lincomycin, vancomycin, tetrazolium violet, and tetrazolium blue. While, some of the tests showed inconsistent readings.

Biochemical features of PPL560^T^ and PPL568 were compared to previously reported strains such as *X. albilineans* CFBP2523^T^, *X. translucens* DSM18974^T^, *X. hyacinthi* CFBP1156^T^, *X. theicola* CFBP4691^T^. All these belong to clade-I of genus *Xanthomonas* along with PPL560^T^ and PPL568 (Trébaol et al., 2000; L Vauterin et al., 1995) Two other rice associated bacterial strains, *X. oryzae* 5047^T^ and *X. maliensis* LMG27592^T^ were also involved in the comparison (Trébaol et al., 2000; Triplett et al., 2015; L Vauterin et al., 1995)**(Table 3)**. Significant differences were observed in the metabolism of α-D-Lactose, D-Meliobiose, β-Methyl-D-Glucoside, L-Fucose, Citric acid, Propionic acid, and acetic acid.

**Table 3:** Phenotypic characterization of PPL560^T^ and PPL568, *X. sontii* PPL1T and *X. sacchari* NCPPB4341T taken as control along with comparison to other related species of genus *Xanthomonas*, LMG27592^T^*, LMG5047^T^*, LMG949^T^*, CFBP2054^T^*, CFBP2054^T^*, CFBP1156^T^*, and CFBP4691^T^ (*data taken from literature). Symbols represent + for source utilization, - for lack of metabolism and v for variable readings.

| BIOLOG tests | PPL560<sup>T</sup> | PPL568 | *X. sontii*<br>PPL1<sup>T</sup> | *X. sacchari*<br>NCPPB4341<sup>T</sup> | *X. maliensis*<br>LMG 27592<sup>T</sup> | *X. oryzae*<br>LMG5047<sup>T</sup> | *X. albilineans*<br>LMG494<sup>T</sup> | *X. translucens*<br>CFBP2054<sup>T</sup> | *X. hyacinthi*<br>CFBP1156<sup>T</sup> | *X. theicola*<br>CFBP4691<sup>T</sup> |
| --- | --- | --- | --- | --- | --- | --- | --- | --- | --- | --- |
| Dextrin | + | + | + | + | + | + | - | + | + | + |
| D-Maltose | + | + | + | + | + | + | - | + | + | + |
| D-Trehalose | + | + | + | + | + | + | - | + | + | + |
| D-Cellobiose | + | + | + | + | + | + | - | + | + | + |
| Gentiobiose | + | + | + | + | + | + | - | - | + | + |
| Sucrose | + | + | + | + | + | + | - | - | + | - |
| D-Turanose | + | + | + | + | v | - | - | - | + | + |
| D-Raffinose | - | - | - | - | - | - | - | - | - | - |
| α-D-Lactose | + | + | + | + | + | - | - | - | - | - |
| D-Melibiose | + | + | + | + | + | - | - | - | - | - |
| β-Methyl-D-Glucoside | + | + | + | + | + | - | - | - | - | - |
| Gelatin | + | + | + | + | + | - | - | - | + | - |
| D-Fructose | + | + | + | + | + | + | + | + | + | + |
| D-Galactose | + | v | + | + | + | - | - | - | + | + |
| L-Fucose | v | + | + | + | + | - | + | - | - | + |
| L-Rhamnose | - | - | - | v | + | - | - | - | - | - |
| D-Sorbitol | - | - | - | - | - | - | - | - | - | - |
| D-Mannitol | - | - | - | - | + | - | - | - | - | - |
| D-Arabitol | - | - | - | - | + | - | - | - | - | - |
| L-Alanine | + | + | + | + | + | + | - | + | - | + |
| L-Aspartic Acid | + | + | + | + | v | - | - | + | + | - |
| L-Glutamic Acid | + | + | + | + | + | + | - | + | + | + |
| Formic Acid | - | v | v | - | v | - | - | - | - | - |
| Quinic Acid | + | v | + | + | - | - | - | - | - | - |
| Citric Acid | + | + | + | + | + | - | - | - | - | - |
| Bromo-Succinic Acid | + | + | + | + | + | + | - | + | + | - |
| Propionic Acid | + | + | + | + | + | - | - | - | - | - |
| Acetic Acid | + | + | + | + | + | - | - | - | - | - |

## Virulence Assay

To test the pathogenicity status of the novel strains we inoculated them on rice, considering the strains were isolated from rice seeds. Briefly, bacterial cultures were grown overnight in PS media (1% peptone and sucrose, pH 7.2) were pelleted and washed with Milli-Q water. Leaves of 60 to 80 days old susceptible rice cultivar Taichung Native 1 (TN1) were clip inoculated with scissors dipped in bacterial suspension adjusted to OD 1 measured at 600nm in Milli-Q water. *X. oryzae* pv. *oryzae* BXO1, a pathogen of rice was taken as a positive control, *X. sontii* PPL1^T^ was taken as a negative control and Milli-Q water was taken as a neutral control. The novel strains, PPL560^T^ and PPL568 did not produce lesions like previously described novel non-pathogenic *Xanthomonas, X. sontii* PPL1^T^ **(Figure 2)**.

**Figure 2.**
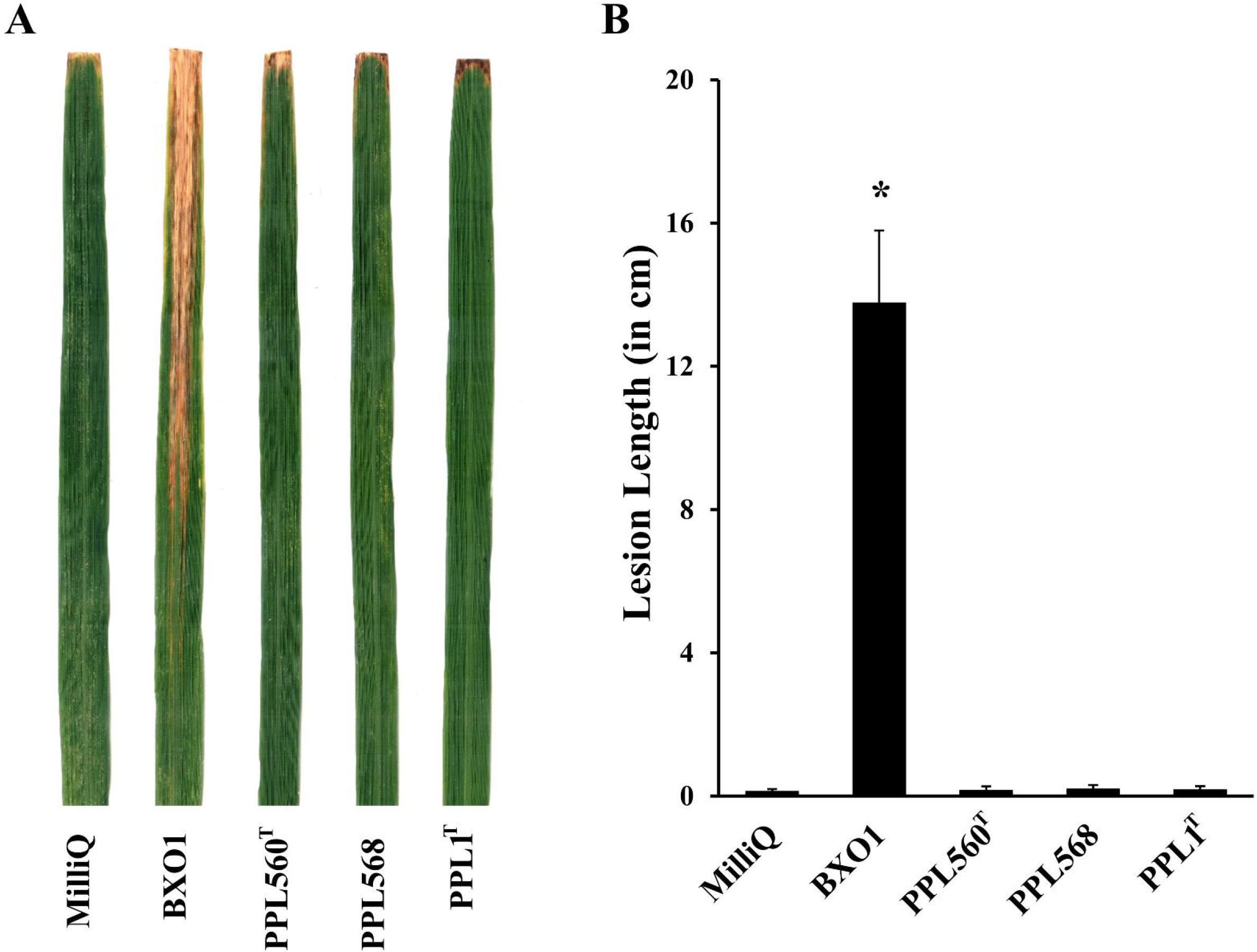
The novel strains do not cause lesions on rice leaves. TN1 rice leaves were clip inoculated with Milli-Q, *X. oryzae* pv. *oryzae* BXO1, novel strains PPL560^T^ and PPL568, and *X. sontii* PPL1^T^. The lesion lengths were tabulated 15 days post-inoculation. **(A)** A sample composite image of infected rice leaves. **(B)** Bar graph of average virulence lesion lengths observed. Error bars used are standard deviations calculated from a minimum of 14 leaves. The * labelling of the column indicates a significant difference in lesion length using unpaired two-tailed student’s *t*-test (*p*-value < 0.01).

## Type III secretion system (T3SS) and effectors (T3Es)

Presence/absence of T3SS (Ryan et al., 2011) and T3Es (http://www.xanthomonas.org) were carried out using tblastn v2.12.0 with cut-off values of query length similarity ≥60% and query sequence identity ≥40%. The T3SS and T3Es profile of PPL560^T^ and PPL568 was compared with their closely related *X. sacchari* CFBP4641^T^, *X. sontii* PPL1^T^ (Bansal et al., 2021) both of which are non-pathogens, and with the pathogen of rice *X. oryzae* pv. *oryzae* strain BXO1 (Kaur, Bansal, Kumar, Sonti, & Patil, 2019). PPL560^T^ and PPL568 lack all the 23 T3Es (XopA, XopAA, XopAB, XopAD, XopAE, XopC2, XopF1, XopI, XopK, XopL, XopN, XopP, XopQ, XopR, XopT, XopU, XopV, XopW, XopX, XopY, XopZ, AvrBs2, and HpaA) similar to PPL1^T^ and CFBP4641^T^. Their T3SS profile is also similar to PPL1^T^ and CFBP4641^T^ where all of these lack T3SS genes except for HrcN **(Figure 3)**. The absence of T3SS genes and T3Es might explain the non-pathogenic behaviour of these strains.

**Figure 3.**
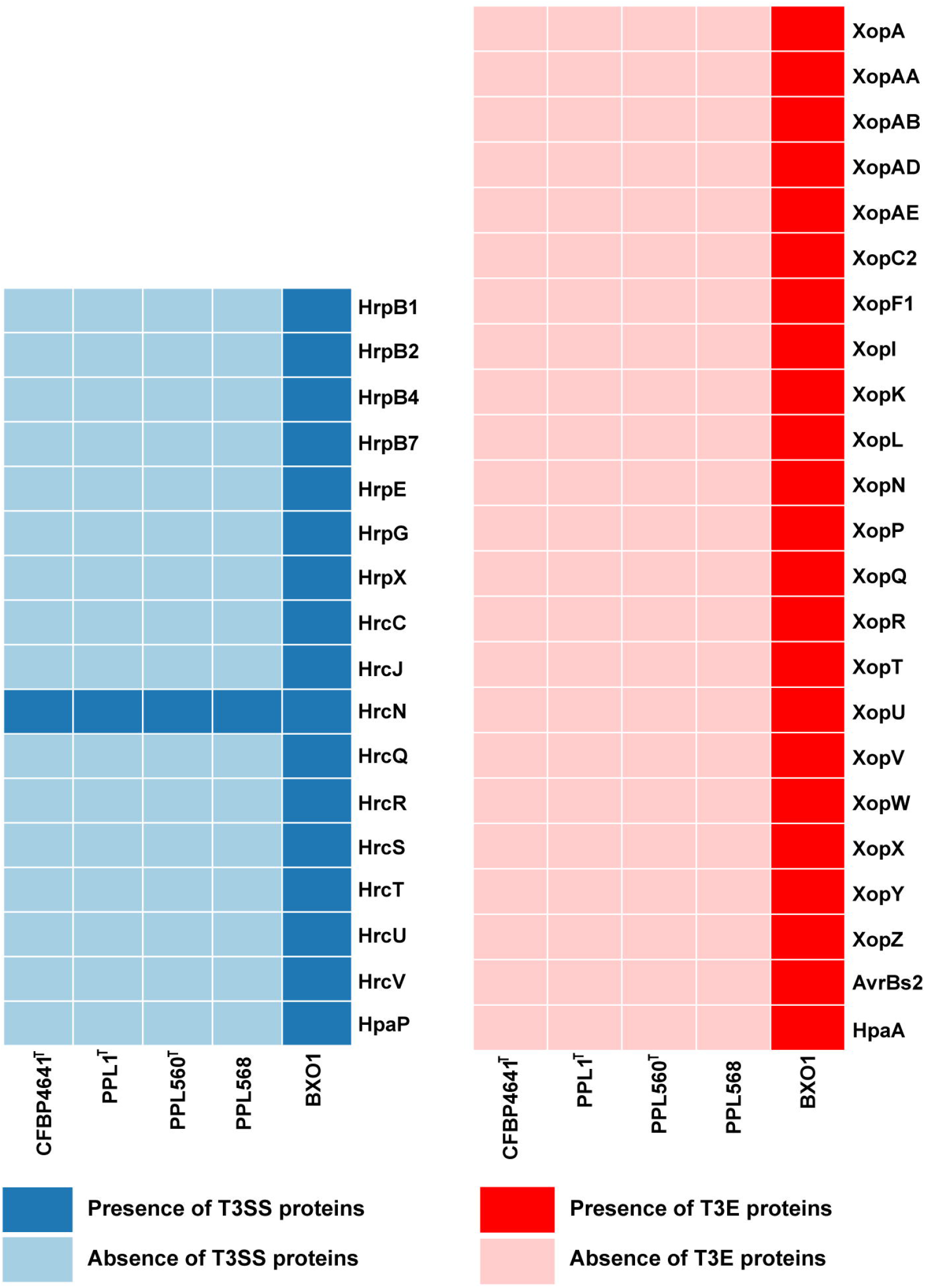
Novel strains lack type III secretion system and its effectors. Presence/Absence profile of genes encoding type III secretion system (T3SS) and type III effectors (T3Es) among PPL560^T^, PPL568, *X. sontii* PPL1^T^, *X. sacchari* CFBP4341^T^ as well as *X. oryzae* BXO1 strains obtained using tBLASTn analysis. The dark and light blue coloured box represents the presence and absence of T3SS genes respectively. Similarly, dark and light red depict the presence and absence of T3Es, respectively.

## Description of *Xanthomonas indica* sp. nov

*Xanthomona indica* (in’di.ca. L. fem. adj. indica indian, named after the country of isolation) Cells isolated from healthy rice seeds are gram-negative, aerobic, and rods shaped. Strains of this species form convex, pale yellow and mucoid colonies after 24-48 hours growth at 28°C on PSA medium. Cells utilizes Dextrin, D-Maltose, D-Trehalose, D-Cellobiose, Gentiobiose, Sucrose, D-Turanose, α-D-Lactose, D-Meliobiose, β-Methyl-DGlucoside, D-salicin, N-Acetyl-D-Glucosamine, α-D-Glucose, D-Mannose, D-Fructose, Gelatin, L-Alanine, L-Aspartic acid, L-Serine, L-Glutamic Acid, Pectin, Methyl pyruvate, L-Lactic acid, Citric acid, L-Malic acid, Bromo-succinic acid, Tween 40, Propionic acid, and Acetic acid. Growth was observed at pH 6.0, 1% NaCl and sodium lactate. Upon leaf clip inoculation, the isolates do not cause any disease symptoms. oANI and dDDH delineates the isolates from reported species. The type strain is PPL560^T^=MTCC13185 and PPL568 is another constituent member of the novel species.

## Abbreviations

OrthoANI: Ortho Average Nucleotide Identity
dDDH: digital DNA-DNA Hybridisation
T3SS: Type III Secretion System
T3Es: Type III Effectors

## Author Contributions

RR isolated the strains, performed identification, and genome analysis. TS did phylogenetic analysis. VNM did the plant virulence assay. RR and AK carried out biochemical tests. RR drafted the manuscript with inputs from KB, AK, TS, RVS, HKP and PBP. PBP planned and participated in the design of the study along with RR, KB, HKP, and RVS. PBP, HKP, and RVS applied for funding and coordinated the study. All authors read and approve the manuscript.

## Acknowledgments

This work is supported by a project entitled “Deciphering the mechanism(s) of host-endophytes co-evolution, enhanced secondary metabolite production and crop productivity. Grant Number: NBRI-IMTECH-MLP48. We acknowledge Council of Scientific and Industrial Research (CSIR) fellowship to RR.

