## Supplementary Figure 1 for "*Xanthomonas indica* sp. nov., a non-pathogenic bacterium isolated from healthy rice *(Oryza sativa)* seeds from India"

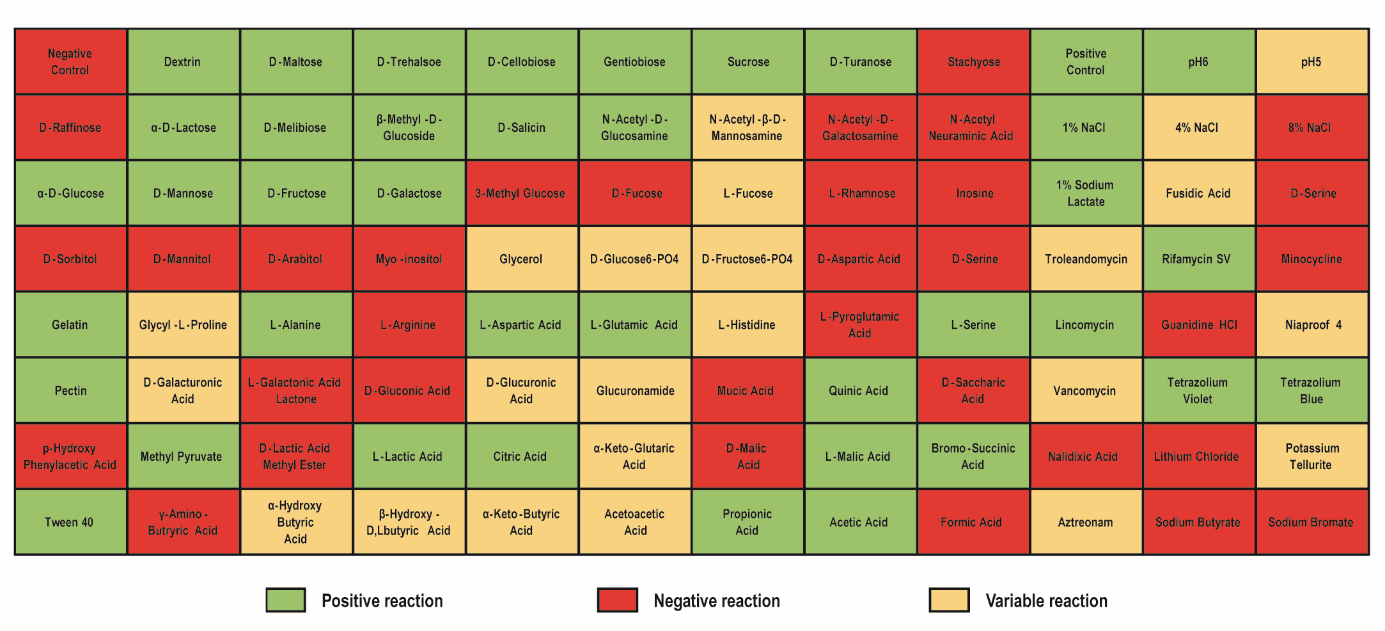

**Supplementary Figure 1:** Biochemical analysis of PPL560^T^ using BIOLOG GN3 microplate showing carbon source utilization, resistance to antibiotics, growth at different pH and salt concentrations. Green, red, and yellow coloured boxes are positive, negative and variable for utilization of their respective sources.
